# REINDEER2: practical abundance index at scale

**DOI:** 10.1101/2025.06.16.659990

**Authors:** Yohan Hernandez-Courbevoie, Mikaël Salson, Chloé Bessière, Hao-liang Xue, Daniel Gautheret, Camille Marchet, Antoine Limasset

## Abstract

Recent advances in biological sequence indexing have enabled the efficient querying of sequence presence across massive genomic data repositories. While presence queries have become tractable at petabyte scale, retrieving quantitative information such as sequence abundances remains a significant algorithmic challenge. Existing abundance-aware indexes are mostly static, difficult to scale, and often trade off completeness, precision, or updatability. We describe a novel discrete abundance index designed for scalability, dynamic updates, and tunable precision. We combine an inverted index with probabilistic and exact structures to support fast, memory-efficient construction and precise high-throughput queries across thousands of RNA datasets.

Our experiments demonstrate that our method REINDEER2 achieves one to two orders of magnitude speedup in construction compared to existing methods, while maintaining comparable or better memory use. Despite using approximate structures for scalability, REINDEER2 achieves sub-1% error on abundance recovery and correlates strongly with reference quantifiers like Kallisto. It also supports sequence-level queries in seconds over thousands of datasets.

**Code and experiments** github.com/Yohan-HernandezCourbevoie/REINDEER2

## 1 Introduction

Significant progress in algorithms and data structures has benefitted the fields of bioinformatics and molecular biology in parallel to progress in DNA and RNA sequencing. The exponential growth of high-throughput sequencing technologies, combined with a dramatic reduction in cost, has enabled data generation overnight in many platforms and hospitals; but poses scalability challenges.

Because data openness is a standard practice in bioinformatics, the reuse of sequencing data from previous studies is encouraged, enabling reanalysis and novel discoveries. Sequencing datasets are massively stored in public repositories, which now hold petabytes of data. Therefore, a fundamental operation is to retrieve in which datasets a query sequence matches. Tools like BLAST, based on pairwise sequence alignment, have been instrumental for such queries. However, their reliance on quadratic-time operations makes them computationally prohibitive at current scales. More recently, the problem has been reframed as data retrieval, and methods have shifted toward the use of fixed-length substrings, known as *k*-mers [17,16]. The core idea behind current strategies is to represent each dataset as a set of *k*-mers and estimate query-dataset similarity through *k*-mer intersections [28], using inverted indexes based on different types of data-structures and compression strategy (e.g. [7,1,20]). These *k*-mer-based approaches have proven scalable and versatile but must scale to billions of *k*-mers and thousands of datasets at least.

Fewer abundance-aware indexing methods extend this principle. In the context of RNA data, querying *k*-mer abundance is as important as detecting presence, since abundance comparison is fundamental to understanding gene regulation [3,4]. Tools such as REINDEER [18], Needle [11], and Metagraph [13] represent the state of the art, enabling exact or approximate *k*-mer abundance queries across massive sequence collections. However, REINDEER remains computationally expensive to build at scale. Metagraph achieves scalability through aggressive *k*-mer filtering, which may miss important signals, especially in cases involving small sequence variants.Entire petabytes-scale databases also exist [8,13]; but while these methods provide a comprehensive overview of our current data, they are resource-intensive to build, making them impractical for new data or clinical settings that need to be built on private servers. Furthermore, their queries remain slow due to the sheer volume of data being scanned. For all these tools, new datasets require complete rebuilding, which is impractical in settings with continuous data inflow. We demonstrate how, using appropriate modeling and data structures, our method can scale to thousands of datasets, support abundance tracking, and allow efficient insertion of new data. Needle introduced a probabilistic query system indexing a subset of *k*-mers with coarse abundance estimates. While it excels as a filter, its approximation strategy may be unsuitable for applications requiring base-level accuracy, such as rare variant detection or complex isoform analysis. In our work, we aim to provide a more precise and tunable approximate scheme that balances scalability with accurate abundance estimation.

## 2 Methods

We introduce the REINDEER2 index, a scalable inverted index for retrieving RNA abundance information across large-scale sequencing datasets. In subection 2.1, we outline the data model and approximations underlying our design. Subsection 2.2 describes strategies for improving scalability, enabling efficient memory usage, and supporting dynamic dataset insertion.

### 2.1 Underlying model

Input representation The primary input to RNA abundance studies consists of sequencing datasets in textual formats composed of four characters or bases, each line representing a fragment of the molecule, called a *read*. To handle the redundancy inherent in sequencing data, where RNA molecules, and *reads*, appear with varying abundance, we first decompose each *read* at all possible positions in substrings of length *k*, called *k*-mers.

It is frequent (and a requirement in many recent indexing methods) to then group *k*-mers into longer superstrings called *unitigs*, which capture contiguous, biologically relevant sequences [6]. Notably, millions of such *unitigs* datasets are readily accessible to the community via the Logan project [9], which removes the burden of *unitig* construction by providing pre-processed, reusable representations of publicly available data. This transformation reduces storage and computational burden at the indexing level, especially because it usually filters out low-abundance, likely-noisy *k*-mers. Previous studies observed that all *k*-mers in a *unitig* have similar abundances [18], therefore standard formats approximate *unitig*-level abundance by averaging their counts, smoothing minor fluctuations at the individual *k*-mer level. More justification to this model can be found in Supplementary. These *unitigs* and their counts are the actual input to REINDEER2. Finally, to reduce index size further, we introduce an novel discretization scheme for abundance values, using a logarithmic encoding with tunable precision (see subsection 2.2).

### *K*-mer set and abundance representation

We use Bloom filters to represent each *k*-mer sets efficiently. The use of discretized abundances enables kmer abundances to be encoded in a matrix-like superposition of Bloom filters, where each row - Bloom filter - is associated with a dataset and an abundance level, as illustrated in Figure 1. When a couple (*k*-mer, abundance) is parsed in a given dataset, it is hashed in the row corresponding to the (dataset, abundance level) row. We detail in subsection 2.2 how abundance levels are computed.

**Fig. 1.**
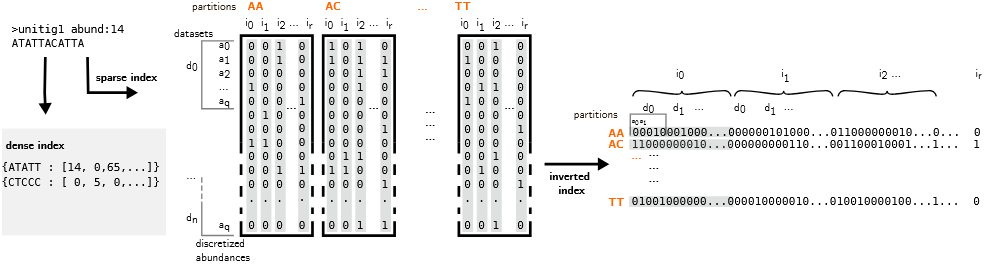
Simplified view of the REINDEER2 index. Left: dense index, where *k*-mer occurring in all or most datasets are stored with explicit and exact abundances. Middle: abstract idea of the sparse structure, where we have one Bloom filter per dataset per abundance (row-wise Bloom filters). Right: actual inverted index of our sparse structure. Each line is a single Bloom filter containing the information of several datasets. Partitioning is showed with lexicographic minimizers of size 2. The first slice of each partition is shown in gray.

As described below; the rationale for using Bloom filters is their support for fast streaming *k*-mer insertion and real-time updates and representing each dataset indepen dently allows easier updates to the index. Following practices from prior works [28,5], we use a single hash function to accelerate queries while controlling the false positive rate through appropriate filter sizing. To minimize space, we encode filters with Roaring bitmaps [15], avoiding explicit storage of all bits.

### Normalization

Sequencing depth in RNA-seq indicates how thoroughly the RNA molecules have been sampled, and can vary between experiments. Therefore, *read* abundances or *k*-mer abundances must be normalized to account for differences in depth across experiments [26]. TPM (Transcripts per million) is a normalization commonly used in RNA studies to account for molecules and sequencing depth abundances when genes are well-known [30] (described in Supplementary).

We work agnostically from known genes, directly on raw data, and with *k*-mers, therefore, we apply a normalization by dividing abundance values by the total number of (non distinct) *k*-mer and using per million as a scaling factor to resemble TPM.

### Inverted index and shared data optimization

REINDEER2 stores abundance data in an inverted index, mapping each *k*-mer to the datasets in which it appears, along with its abundance. This structure allows direct access to abundance values across datasets with a single random access, rather than querying each dataset separately. A naive per-dataset *k*-mer index leads to redundant storage of shared sequences, especially common in RNA-seq from the same species (e.g., housekeeping genes). In contrast to prior literature, REINDEER2 instead separates *k*-mers into two categories: *dense k-mers* found in most datasets, stored in an exact index, and *sparse k-mers* found in fewer datasets, stored in Bloom filters. This hybrid design reduces Bloom filter load, improving accuracy and reducing false positives by decreasing the Bloom filters load factor. Exact *k*-mers are identified on-the-fly during construction (details in subsection 2.2).

### Query model and output

A typical query consists of a *read* or RNA molecule (often >100 bases). Queries are decomposed into overlapping *k*-mers, which are intersected with the index, and resuts are reported if the intersection size is larger than a threshold (default 50% of the query set). This threshold is important to account for *k*-mer content that may differ due to errors or small mutations. For each dataset, the median abundance across all matching *k*-mers of the query is returned (see Figure 3) in a csv file where rows are (query, abundance, dataset) triplets.

Two types of possible errors can be described. The first type occurs when a collision happens in the dataset that is screened for abundance, leading to several possible abundance levels. We keep the minimum, therefore the abundance values can be underestimated in some cases. The second type is a bit occurring in a dataset in which the *k*-mer was not in the first place, thus we report an abundance despite the *k*-mer absence (see Figure 3 right). Importantly, false positives from Bloom filters are mitigated by query length: the likelihood that all *k*-mers in a query are false positives decreases geometrically with the number of *k*-mers [28,5]. We analyze query performance and accuracy in subsections 3.4 and 3.6.

### 2.2 Strategies for improved scalability

#### Bloom filter factorization

To mitigate the significant memory overhead from using up to billions of Roaring bitmaps, we encode multiple data slices into a single, larger bitmap. This is achieved in a collision-free manner by assigning a unique offset to the identifiers within each slice. This strategy effectively amortizes the storage overhead across the combined slices while still permitting independent queries on each original slice within the aggregated bitmap, as illustrated in Figure 2.

**Fig. 2.**
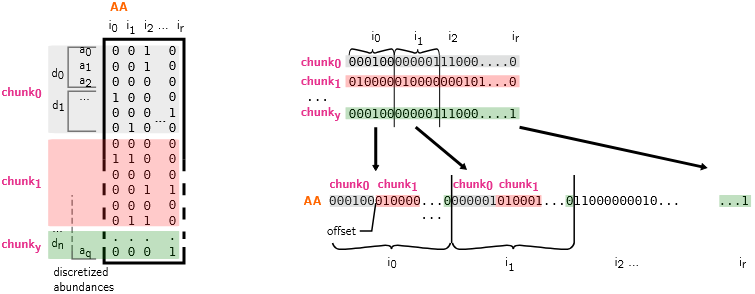
Schematic view of the sparse index division and merge. The example is showed on a single partition for simplicity. On the left, an abstract view of the structure divided in chunks. On the right, bits of each chunks are added to a single Bloom filter per partition, by doing a union of chunks with offsets.

**Fig. 3.**
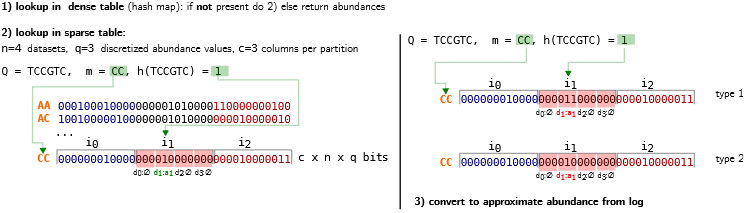
Examples of query at the *k*-mer level. Left: query mechanism in REINDEER2. If it is not found in the dense index, queried *k*-mers are hashed to find their indices in the filter and their partitions are determined by minimizers. Then all datasets are screened to find bits indicating abundances (here abundance level 1 in dataset 1). Right: two types of possible false positives. The first type can lead to an under estimation of the abundance level when different abundance levels are reported for a given dataset. The second type is a true false positive, reporting an abundance in a dataset where the *k*-mer is absent.

#### Partitioning

Minimizers are the smallest *m*-mer with *m<k* in a given *k*-mer, according to a defined order [24]. Minimizer-based partitioning has demonstrated its efficiency for *k*-mer indexing [20,12,19]; therefore, we assign *k*-mers to partitions based on their minimizer rather than at random (Figure 1 shows a simplified example with lexicographic minimizers). This strategy leverages the observation that consecutive *k*-mers often share the same minimizer, which enables buffered operations that improve both construction and query times by optimizing cache coherence and allow coarse-grained parallelism [23]. Indeed, using hundreds or thousands of partitions reduces the size of the loaded substructure by orders of magnitude, both during construction and at query time. This allows the structure to utilize lower-level caches, thereby accelerating performance. Although the distribution of minimizers in genomic data is known to be skewed [21], we uniformly shuffle a large set of minimizers within each partition to mitigate this bias, resulting in relatively uniform partition sizes in practice.

#### Merge and update of REINDEER2 indexes

Indexing billions of k-mers from very large collections can cause a peak in RAM usage that exceeds available memory, even when using compressed structures. To manage this, REINDEER2 rely on its partition system to builds multiple sub-indexes sequentially, effectively lowering this memory peak. Once constructed, these sub-indexes are merged into a final, global index. This merging process is also memory-efficient as it operates on a partition-by-partition basis. At the partition level, merging involves a simple bitmap remapping. For each sub-index, local column positions are converted to global ones by adding an offset. All these remapped bits are then combined into the final, unified bitmap for that partition.

Furthermore, this merging mechanism makes the index inherently updatable. A new dataset can be indexed separately and then efficiently merged with the main index, enabling seamless, incremental updates.

#### Logarithmic quantization scheme for abundances

One distinctive feature of Needle is its ability to compute optimal quantization levels based on the observed distribution within the indexed collection. However, this approach is computationally expensive and does not scale well when a large number of levels, necessary for high precision, is required. Moreover, maintaining these optimal levels becomes non-trivial when incorporating novel datasets. Here, we propose a more general approach that leverages logarithmic scales to achieve tunable precision over an arbitrarily large input range. Once the maximum abundance is specified, our method guarantees that the quantization system remains fixed even as new datasets are added, thereby enabling efficient updates.

REINDEER introduced the idea of approximating the abundances with the base-2 logarithm each abundance. However, this has the drawback of being a rough approximation. We adopt a strategy based on this idea, but adapting the logarithmic base to achieve a tunable precision.

To determine the base that provides *Q* levels for handling abundances up to *Max_Abundance*, we start by considering the formula: *Q*≤log_*B*_(*Max_Abundance*)*< Q*+1. Note that this quantization is often suboptimal because certain abundance values are not observed in practice, *ie*. when ⌊log_*B*_(*q*)⌋+1*<*⌊log_*B*_(*q*+1)⌋, with *q* ∈ ℕ^∗^.

To address this issue where some values would not be used, we employ an identity function for lower abundances while applying logarithmic scaling to higher values. The threshold *t*, at which we switch from the identity function to logarithmic scaling, is defined as follows 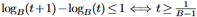, for *B >* 1. Converting an abundance *x* to its log-transformed value is achieved with:

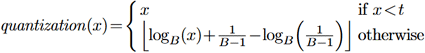

Using this function, one can determine the optimal base *B*, given the maximal abundance (*Max_Abundance*) and the number of abundance levels *Q*. We provide all the details on the quantization in the Appendix.

In practice, as shown in Figure 4 (left), we observe that using 256 abundance levels allows us to achieve high precision (near 1%), and that the overall abundance range to be handled has a significant impact on the final precision.

**Fig. 4.**
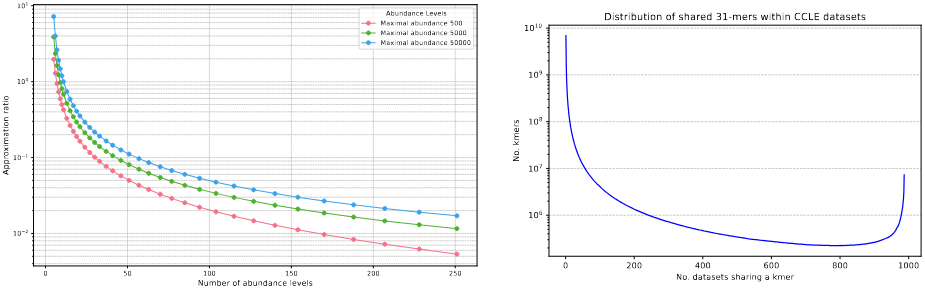
Left: abundance precision respective to the amount of abundance levels used and the maximal abundance handled. For the maximal number of abundance levels (256), the base *B* is 1.010 for *Max_Abundance*=500, 1.023 for *Max_Abundance*=5,000 and 1.033 for *Max_Abundance*=50,000. Right: distribution of 31-mers according to the number of CCLE datasets in which they appear.

#### Handling of dense *k*-mers

Inserting ubiquitous *k*-mers into thousands of filters is suboptimal with respect to indexing time, index size, and query efficiency. As illustrated in Figure 4 (right) using a human cancer sample collection (CCLE, detailed in subsection 3.1), the prevalence of commonly shared *k*-mers justifies a specialized handling strategy. Notably, this approach is optional and is particularly beneficial when a significant portion of the dataset content is highly shared.

To address this issue, we propose to handle *dense k-mers*, those appearing in most datasets, separately from *sparse k-mers*, which occur in fewer datasets. In an online procedure, sparse *k*-mers are inserted into the primary index, while dense *k*-mers are stored in a dedicated hash table that maps each dense *k*-mer to its abundance vector. At regular intervals, we re-evaluate the dense *k*-mers to ensure they continue to satisfy the density criterion by iterating on the abundance vectors. If a *k*-mer no longer qualifies as dense, it is removed from the dense table and reinserted into the sparse index according to its occurrences and abundances. This strategy avoids redundant insertions of *k*-mers common to thousands of datasets and enables more efficient storage of their abundance vectors.

At query time, each *k*-mer is first searched for in the dense table; if it is not found there, the search proceeds to the sparse index. In practice, this strategy incurs negligible overhead in terms of query time and index size, while enhancing precision via two mechanisms. First, dense *k*-mers are stored exactly, eliminating the possibility of false positives. Second, by excluding dense *k*-mers from the Bloom filters in the sparse index, the overall false positive rate for sparse *k*-mers is reduced.

## 3 Results

### 3.1 Datasets and sequences description

We used pre-computed *unitigs* from Logan Project [9] constructed from the Cancer Cell Line Encyclopedia (CCLE) collection [2] that includes curated sequencing data from more than 1,000 different human cancer lab-grown cells. We used *unitigs* from the Logan initiative, that provides cleaned *unitigs* of 987 of the CCLE datasets, built for *k*=31 for any *k*-mer seen more than twice. Therefore all our indexes are built for the same *k* value, and very rare *k*-mers may be absent. The whole collection represents a rough 11.6 billion *unitigs*,with 164 billion 31-mers (statistics on sequences were computed with seqkit [27]). In addition, we selected a second dataset to demonstrate our scalability capabilities. We filtered and downloaded 10,000 human RNA-seq accessions from the European Nucleotide Archive so that the datasets contain at least 10 million *reads*, and are associated with melanoma/melanocyte meta-data. The corresponding *unitigs* represent a total of 67 billion sequences and 3.02 trillion bases or 992 billion 31-mers.

### 3.2 REINDEER2 outperforms state-of-the art in resource usage for construction

First, we wanted to describe the inherent advantages and drawbacks of using state of-the-art abundance indexes on a server with 128GB of RAM, namely REINDEER, Needle, and Metagraph, in comparison to REINDEER2. We excluded recent structures dedicated to indexing abundances of a single dataset, such as [22].

We constructed indexes representing subsets of the CCLE collections with growing numbers of datasets. We reported the results in Figure 5. We observed that REINDEER2 is one or several orders of magnitude faster to construct than all its competitors. It is also memory-efficient, thanks to its merging system, which prevents high memory usage peaks. While index size optimization is not the primary focus, the resulting index remains practical and can even be competitive with state of-the-art approaches, except for Needle achieving smaller index sizes because of sampling.

**Fig. 5.**
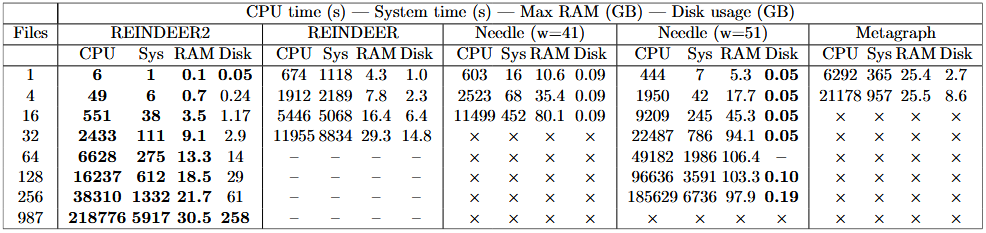
Resource usage during sparse index construction on CCLE collection. “*×*” stands for tools that failed to run and “–” for tools that were not run by lack of time.

It is worth noting that this benchmark is not entirely fair for several reasons. Needle indexes only a subset of *k*-mers with false positives, Metagraph performs stringent graph cleaning before indexing, while REINDEER indexes all *k*-mers of *unitigs* exactly. More importantly, REINDEER2, like its predecessor, relies on pre-constructed *unitigs*, which can be orders of magnitude smaller than the original FASTQ files used by Needle or Metagraph. However, the pre-processing steps of Metagraph and Needle cannot be isolated from their own usage, making a direct comparison challenging.

We still argue that this benchmark effectively highlights the practical usage cost of these tools, as *unitigs* are emerging as efficient alternatives to FASTQ files and extremely efficient techniques now exist to compute them rapidly and in a memory-efficient manner [10,14] (typically around 1 hour for a whole genome human dataset, usually larger than RNA samples) when they are not already available [8].

### 3.3 Increased precision for a marginal overhead

In the previous section, we used a low number of quantization levels (16) to align with the capabilities of Needle and the original REINDEER in logarithm base 2. In REINDEER2, however, we advocate for high-precision abundance quantization, which involves using a substantially larger number of levels. A natural concern is that this increased precision might significantly increase the cost of both index construction and query operations. Similarly, employing larger Bloom filters to achieve lower false positive rates could result in higher resource consumption, potentially limiting the method’s practicality in some applications. To evaluate these concerns, we experimented with various settings for the number of abundance levels and a wide range of Bloom filter sizes (from 2^26^≈67M bits to 2^34^≈17G bits). Figure 6 reports the corresponding construction times and index sizes. Despite exponential increases in both Bloom filter size and abundance levels, resulting in the number of implicit bits increasing by more than four orders of magnitude, the overall index sizes increases by less than an order of magnitude. This highlights the efficiency of the sparse data structure relying on Roaring and Bloom filters factorization, and the fact that high precision indexing is affordable in practice.

**Fig. 6.**
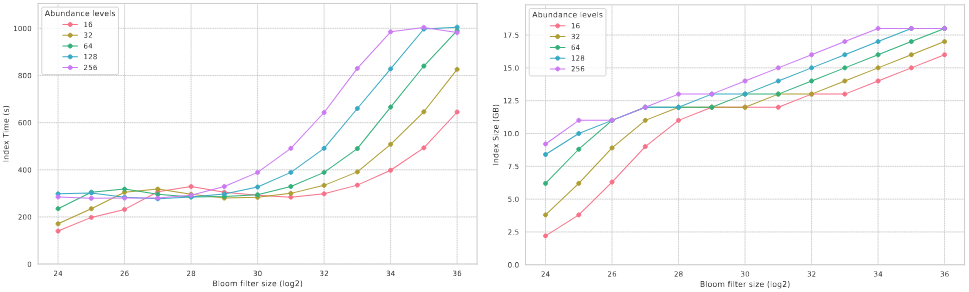
Construction time (left) and index size (right) according to the number of abundance levels on various Bloom filter size of 2^*b*^on 128 files from CCLE collection (sparse index).

### 3.4 High throughput queries

Query performances are known to degrade as indexes go larger. Here we wanted to assess the impact of index size, driven both by Bloom filter size and number of abundance levels, on query performances (Figure 7). We queried human full-length RNAs (10,851 Refseq human transcripts) as positive queries, and sequences unrelated to the index content (randomly generated), as negative queries to have a better idea of “real life” performance. In a first experiment we increased Bloom filter sizes and abundances levels on 128 files of the CCLE collection. We queried 10,851 human transcripts to observe that the bloom filter size has little impact on query throughput, in contrast to the number of abundance levels. In a second experiment, for the same query, we selected a large Bloom filter size (2^32^) and a high number of abundance levels (256).

**Fig. 7.**
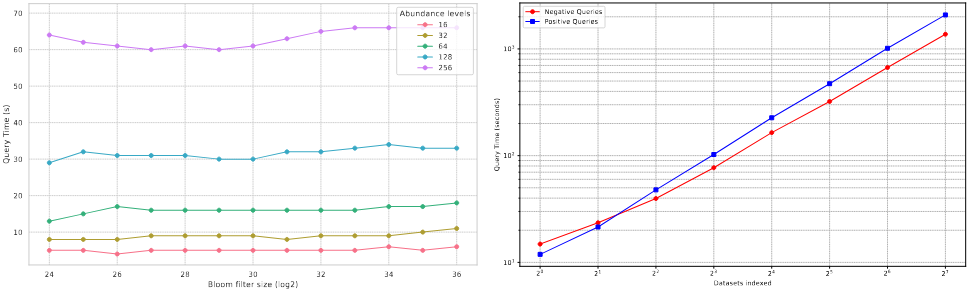
Query time in the sparse index, according to the number of abundance levels on various Bloom filter size of 2^*b*^on 128 files from CCLE collection (left). Time to perform a batch of queries on a growing index size (right). The positives queries are 10,851 human transcripts totalling 40 millions bases. The negatives queries are a file with matching number of sequences and length distribution composed of random sequences.

The index showed high efficiency, processing millions of checks per second. As expected, the running time increases linearly with the number of documents in the index, with negative queries being noticeably faster to execute. Since each query must be evaluated across all indexed datasets, the effective query throughput is inversely proportional to the number of documents in the index.

In order to compare REINDEER2’s performance with that of its competitors, we queried CCLE indexes that could be constructed for all methods; with increasing batches sequences of growing sizes. The results presented in Supplementary Figure S1 show that the improvements made to the construction phase did not come at the expense of query time, REINDEER2 remains comparable to its direct competitors and notably outperforms REINDEER.

### 3.5 Efficient scalability on RNA-seq datasets

To assess the scalability of REINDEER2, we indexed progressively larger collections of melanoma datasets, increasing both the number and size of the files. The first collection, comprising one thousand small files, was indexed in under 40 minutes, processing 0.48 billion sequences and 30 billion bases. The second collection, which included two thousand medium-sized files, required 100 minutes to index 1.25 billion sequences and 76 billion bases. Indexing the third collection, consisting of five thousand large files, took 10 hours to process 6.42 billion sequences and 342 billion bases. Finally, we indexed the entire dataset, including the largest files. This final step, covering a total of 67 billion sequences and 3.02 trillion bases, was completed in 2 days and 10 hours using 64 threads.

### 3.6 Cross-tool quantification accuracy of REINDEER2

Here we wanted to validate that REINDEER2’s abundance estimations could be trusted for research experiments. We compared REINDEER2’s results to REINDEER to assess the impact of false positives in comparison to an exact structure. A complete review of REINDEER’s quantification accuracy was also performed in [3]), showing REINDEER yields results well correlated even to *in vitro* precise quantification techniques, therefore it can serve as a direct mean of comparison for our approach. We also added Kallisto [29] to the comparison, that is one of the most established RNA quantification tool, designed to work at the single dataset level and using well known RNA molecule models. Kallisto’s index is built on a fundamentally different input, making direct performance comparisons with our method inappropriate. We selected a subset of 10 CCLE datasets where 1,000 human genes were quantified using both Kallisto and REINDEER (settings are provided in Appendix). REINDEER2 was used to query directly the CCLE indexes built in the previous sections, with 255 abundance levels and a Bloom filter size of 2^32^. We expected differences between approaches, since Kallisto assigns sequences to the different molecules of the database based on an expectation-maximization algorithm to resolve multiple assignments. Kallisto can yield raw counts and normalized counts using TPM (see subsection 2.1). REINDEER and REINDEER2’s result must be closer since they index the samples and quantify a query sequence by seeking its *k*-mers in the index, and their normalizations are the same.

REINDEER2 showed strong correlation with the exact index of REINDEER. In order to separate the effect of the median of *k*-mer counts returned by REINDEER2 for query sequences larger than *k*, we verified the correlation between REINDEER and REINDEER2 at the single *k*-mer level (Figure 8 top row). It is worth noting that handling dense *k*-mers separately enhances accuracy since their abundances are stored exactly and this reduces the overall false positives of the sparse index. This strategy improves results compared to a sparse-only insertion approach, as illustrated in Figure 8, top row, right. Query sequences longer than 1,000 bases were less impacted by false positives, leading to improved correlations for the gene queries compared to *k*-mer queries (middle row, right with dense *k*-mers handling). The correlation with Kallisto confirms that REINDEER2 is in the same ballpark as REINDEER (based on [3]’s results) when comparing Kallisto’s results, especially for normalized counts. In all experiments, querying the sequences corresponding to 1,000 genes took approximately 6 to 8 seconds.

**Fig. 8.**
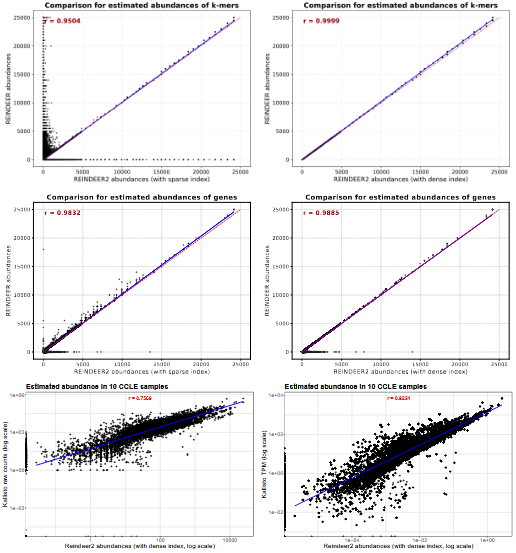
Pearson correlations for quantifying 1000 genes in a CCLE subset. In the top row, REINDEER and REINDEER2 are compared for queries at the *k*-mer level, with top right figure showing REINDEER2’s regular strategy where dense *k*-mers are handled outside of Bloom filters and top left figure shows the impact on the query accuracy when these dense *k*-mers are inserted in filters. Middle row shows similar comparison for “real life” queries at the gene level. Bottom row shows comparisons to Kallisto’s TPM and raw counts with REINDEER2’s regular index. Bottom figures are shown in log-log scale to better visualize small and medium values.

## 4 Discussion

We presented REINDEER2, a scalable index enabling fast, accurate *k*-mer abundance queries across large datasets via an inverted structure with tunable precision and dense *k*-mer handling. Compared to prior methods, REINDEER2 offers superior speed, dynamicity, and maintains strong agreement with established quantifiers like Kallisto. Our design benefits from Bloom filter factorization and partitioning. While current probabilistic structures don’t fully exploit *k*-mer consecutiveness, future improvements could reduce redundancy and enhance compression. Abundance estimates are robust overall, though short-sequence queries and high-multiplicity *k*-mers remain challenging. Alternative aggregation and collision-handling strategies (e.g. [25]) may improve accuracy further. Finally, our discretization scheme opens paths for richer metadata indexing and dynamic updates, supporting future extensions to transactional and real-time operations on large, evolving datasets.

## Supplementary material for REINDEER2: practical abundance index at scale

### S1 Methods

#### S1.1 Unitig model for biological information

First, to account for DNA strand ambiguity introduced during library preparation, we convert each *k*-mer to its *canonical* form, i.e., the lexicographically smaller of the forward and reverse complement, that allows to find *k*-mer from both strands during query.

In RNA sequencing, the number of *reads* covering a given position in the transcriptome varies mainly due to two factors: (1) the true abundance of the RNA molecule at that position, which reflects gene expression levels and biological variation; and (2) the sequencing depth, which indicates how many times the RNA was sampled and replicated to ensure sufficient coverage of all positions.

*K*-mer abundances are a proxy of *read* abundances. When grouping *k*-mers into unitigs, our working hypothesis is to expect that (1) the true *k*-mer abundances across the unitig are very similar, and (2) that the coverage of this region is uniform. (2) is known to be imperfectly true for the Illumina sequencing technology [2], and remains a reasonable hypothesis in a majority of bioinformatics methods.

(1) works because of the adequation between unitigs and biological variants. Unitigs are constructed from a De Bruijn graph, where each observed *k*-mer in the dataset is a node, and edges connect *k*-mers that overlap by *k−*1 bases. A unitig corresponds to a maximal path in this graph where each node has exactly one incoming and one outgoing edge, except at the ends.

Branching in the graph, where a node has multiple incoming or outgoing edges, is typically caused by sequence variants. In RNA, a common source of such variation is alternative splicing, where different transcript forms from the same gene reuse shared segments in various combinations. These forms often have distinct expression levels, which result in different abundances for the *k*-mers they contain.

Importantly, such branching patterns introduce breaks in the graph that define the boundaries of unitigs. As a result, each unitig generally corresponds to a sequence region with consistent abundance, since variation tends to occur at unitig junctions (see e.g. [3]). When abundance shifts are observed within a region, they often correlate with a switch from one unitig to another due to underlying structural variation, which justifies our approach.

#### S1.2 Description of Transcript Per Million

In the case genes are known and well described, methods calculate TPM (Transcripts Per Million - transcripts are full length RNA molecules). For TPM we first divide the *read* count for each gene by the gene’s length in kilobases to get reads per kilobase. Then, we sum all the read per kilobase values in the sample and divide each gene’s read per kilobase value by this total (scaled to millions). This final value is the TPM. By normalizing for gene length first and sequencing depth second, TPM provides expression values that are comparable across both genes and samples.

#### S1.3 Quantization

The threshold *t* that determines the limit between the identity function and the logarithmic quantization is computed with:

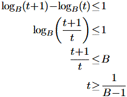

As described in the main text, the quantization of abundance *x* is obtained using the formula below:

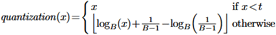

In the above equation, the term 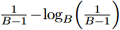 acts as a shift, compensating for the number of values that are exactly represented by the identity function. Using this equation, we determine the optimal base *B*, given the maximal abundance (*Max_Abundance*) and the number of abundance levels *Q*. We require that

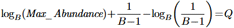

This function is monotonically decreasing for 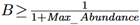, so the root of

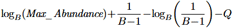

can be found using the bisection method, which yields an approximation for *B*.

### S2 Experiments

#### S2.1 Query performances benchmark

Table S1 presents the query times with batches of 1 to 1,000 gene sequences for the different indexes built on 4 datasets. As expected Needle is faster as it indexes only a fraction of the input *k*-mers but REINDEER2 outperforms REINDEER and is also faster than Metagraph.

**Fig. S1.**
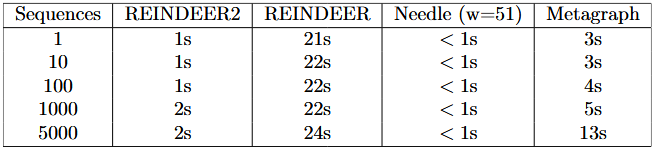
Query time on an increasing number of sequences with an average size of 591 bases. The indexes used are those constructed from 4 datasets on the construction benchmarks presented in the main text. As with the other tools, Needle times - ranging from 0.05s to 0.88s - are rounded to the second.

#### S2.2 Quantification accuracy

Kallisto was run using the Ensembl human reference v108 annotation, which provides a standardized and comprehensive set of gene and transcript definitions for the human genome. REINDEER was built on the Logan unitigs of the 10 CCLE samples and queried using default parameters for indexation and retrieval (*k*=31). For raw counts and transcripts per million (TPM) that were extracted we kept similar settings to those in [1].

